# Direct carbon capture for production of high-performance biodegradable plastic by cyanobacterial cell factory

**DOI:** 10.1101/2021.10.04.462501

**Authors:** Chunlin Tan, Fei Tao, Ping Xu

## Abstract

Plastic pollution has become one of the most pressing environmental issues today, leading to an urgent need to develop biodegradable plastics^1-3^. Polylactic acid (PLA) is one of the most promising biodegradable materials because of its potential applications in disposable packaging, agriculture, medicine, and printing filaments for 3D printers^4-6^. However, current biosynthesis of PLA entirely uses edible biomass as feedstock, which leads to competition for resources between material production and food supply^7,8^. Meanwhile, excessive emission of CO_2_ that is the most abundant carbon source aggravates global warming, and climate instability. Herein, we first developed a cyanobacterial cell factory for the *de novo* biosynthesis of PLA directly from CO_2_, using a combinational strategy of metabolic engineering and high-density cultivation (HDC). Firstly, the heterologous pathway for PLA production, which involves engineered D-lactic dehydrogenase (LDH), propionate CoA-transferase (PCT), and polyhydroxyalkanoate (PHA) synthase, was introduced into *Synechococcus elongatus* PCC7942. Subsequently, different metabolic engineering strategies, including pathway debottlenecking, acetyl-CoA self-circulation, and carbon-flux redirection, were systematically applied, resulting in approximately 19-fold increase to 15 mg/g dry cell weight (DCW) PLA compared to the control. In addition, HDC increased cell density by 10-fold. Finally, the PLA titer of 108 mg/L (corresponding to 23 mg/g DCW) was obtained, approximately 270 times higher than that obtained from the initially constructed strain. Moreover, molecular weight (M_w_, 62.5 kDa; M_n_, 32.8 kDa) of PLA produced by this strategy was among the highest reported levels. This study sheds a bright light on the prospects of plastic production from CO_2_ using cyanobacterial cell factories.

## Main

Increasing CO_2_ concentration in the atmosphere is a severe problem that aggravates global warming and climate instability. Carbon capture has been considered as one of the top priorities for reducing CO_2_ emissions. In addition, plastic pollution has become another pressing environmental issue, as the resilience against degradation and enormous threat to global ecology. Therefore, it is also an urgent need to develop low-cost biodegradable plastics such as polylactic acid (PLA), which serve as the primary way for alleviating plastic pollution. Here, we developed a cyanobacterial cell factory to produce PLA directly from CO_2_, providing a promising method to simultaneously address plastic pollution and excessive CO_2_ emissions.

### Engineering of cyanobacterial cell factories for production of PLA

PLA is a widely used biodegradable material, mainly produced from polymerization of lactic acid or lactide ring-opening polymerization *via* a complicated chemical process ^6^. Herein, an autotrophic microbial cell factory was first constructed for *de novo* biosynthesis of PLA from CO_2_ (the 3^rd^ generation biorefineries ^9^), presented in Fig. 1.

**Fig. 1.**
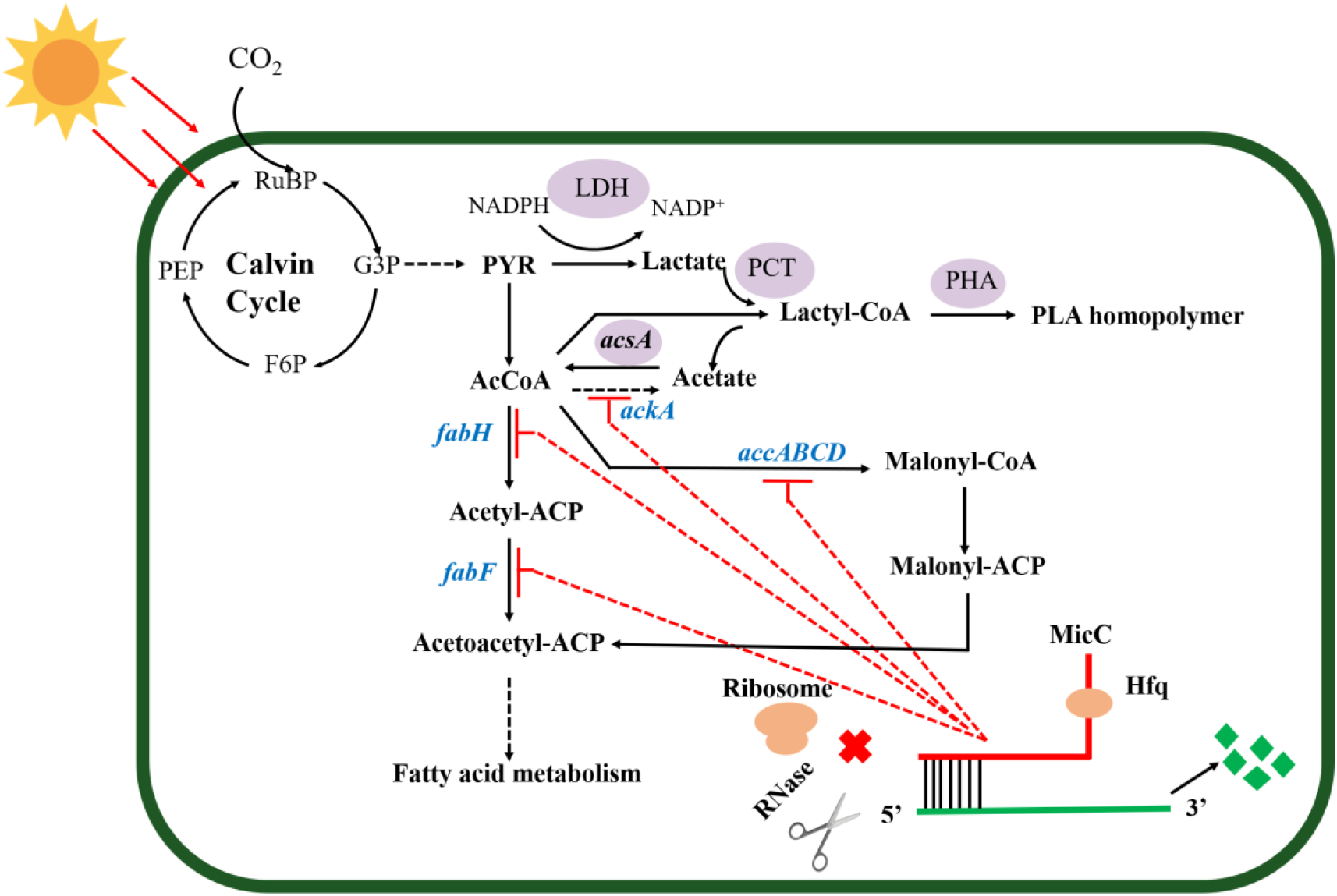
Synthetic pathway for *de novo* biosynthesis of PLA from CO_2_ in autotrophic *S. elongatus* PCC7942. The heterologous expression enzymes which participate in the PLA biosynthesis pathway are shaded in purple. The color blue indicates the essential enzymes for fatty acid metabolism. The Hfq-MicC tools were introduced to the chassis to downregulate the critical genes for redirecting the carbon flux from fatty acid to PLA biosynthesis. Abbreviations: PEP, phosphoenolpyruvate; G3P, glyceraldehyde–3–phosphate; PYR, pyruvic acid; F6P: Fructose 6-phosphate; RuBP, Ribulose-1,5-bisphosphate; AcCoA, acetyl-CoA.

The cassette of biosynthesis for PLA consists of D-lactate dehydrogenase (LDH, encoded by *ldhD*), propionate CoA-transferase (PCT, encoded by *pct*), and PHA synthase (PHA, encoded by *pha*). Cyanobacteria produce NADPH as the primary carrier for reducing equivalents ^7^. Then, LDH was engineered to switch the coenzyme specificity from NADH to NADPH, thus making it more efficient for producing D-lactate (this work was done previously ^7^). Subsequently, the engineered *ldhD*, the evolved *pct*, and *pha*, which come from *Lactobacillus bulgaricus* ^7^, *propionicum*, and *Pseudomonas* sp ^10^, respectively, were cloned into the plasmid pAM-MCS12 under the control of the inducible promoter, P_trc_. The resulting plasmid pAM-PYLW01 was used for recombination at the neutral site I (NSI) of the *S. elongatus* chromosomes (Fig. 2A, 2C). Correct chromosome integration was confirmed by colony PCR (Fig. S1) and DNA sequencing.

**Fig. 2.**
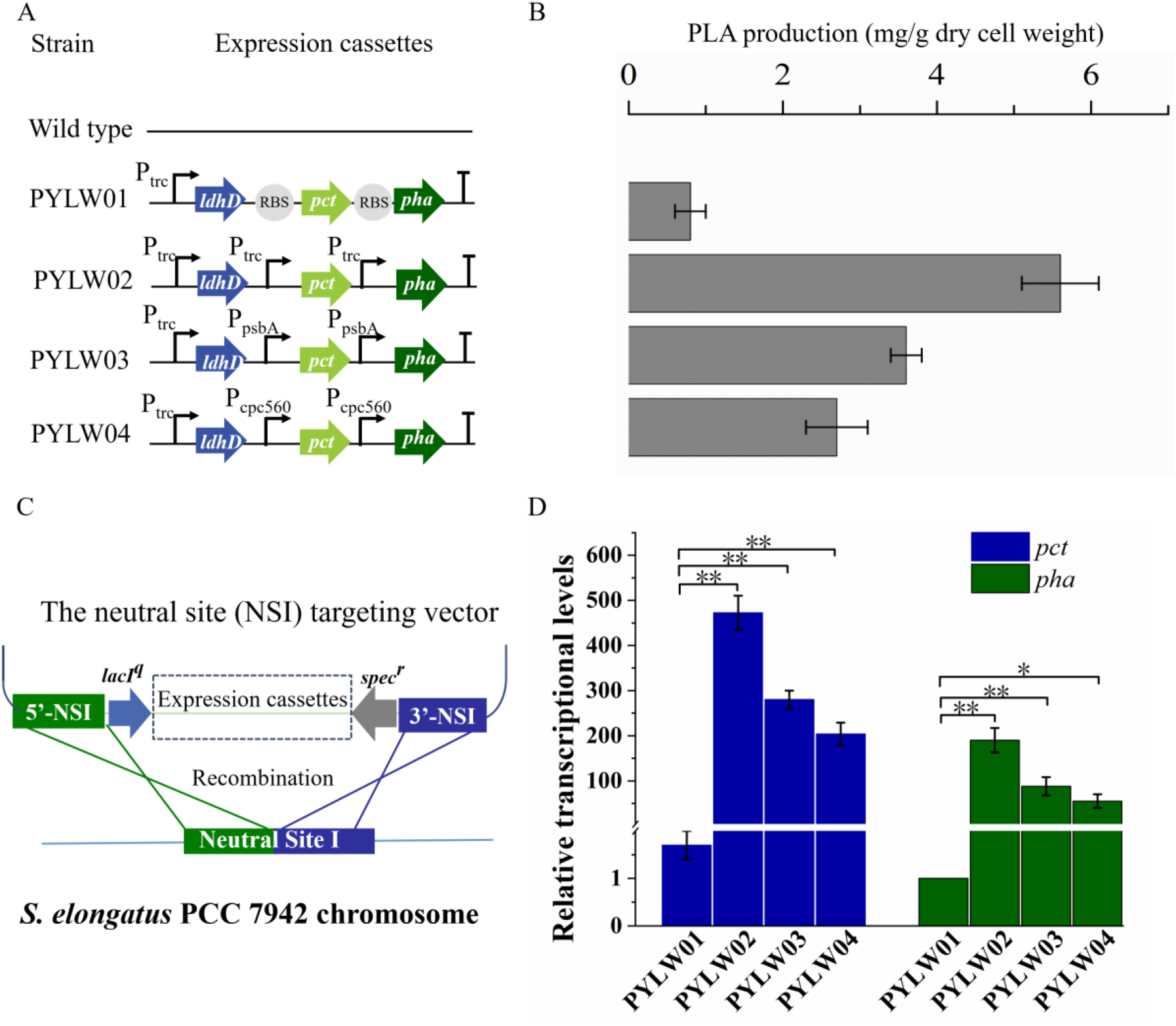
The promoter optimization for the PCT and PHA expression. (A) The construction of different expression cassettes for the PCT and PHA. (B) The PLA production using different expression cassettes. (C) Schematic representation of the expression cassettes recombination into the neutral site I (NSI) of the *S. elongatus* PCC7942 chromosome. *spec*^*r*^ (spectinomycin resistance gene), P_trc_ (promoter trc), P_psbA_ (promoter psbA), P_cpc560_ (promoter cpc560). (D) Quantitative reverse– transcription PCR analysis of relative transcriptional levels of the gene *pct* and *pha*. The housekeeping gene *rnpB* normalized all data (^*^means p-value ≤ 0.05, and ^**^ means p-value ≤ 0. 001). Error bars indicate SD (n = 3)

### Characterization of PLA synthesized by engineered *S. elongatus*

To characterize the PLA granules synthesized by engineered *S. elongatus*, we conducted the transmission electron microscopy analysis for the transformants of strain PYLW07. The PLA polymers accumulated as inclusion granules in engineered *S. elongatus* cells (bottom left and right panel of Fig. 4A), were significantly different from the wild type (top left panel of Fig. 4A). This is in accordance with a previous study ^11^. The 1D (^1^H and ^13^C) and 2D (^1^H-^1^H) COSY NMR spectroscopies were performed to test further the composition and the monomer sequence of the PLA polymer. As shown in Fig 4B, the 600MHz ^1^H spectrum displays the specific quadruplet of PLA homopolymers; the oxymethine proton (-OCH-) and the methyl protons (-CH_3_) of PLA are assigned at 5.2 and 1.6 ppm in the 600-MHz ^1^H spectrum, respectively. Moreover, the ^1^H-^1^H COSY spectrum shows that proton signals at 5.2 ppm (-OCH-) and 1.6 ppm (-CH_3_) are coupled to each other, while other peaks are not associated with a different one (Fig. 4C). In the ^13^C NMR spectrum, the carbonyl carbon (-OCO-), oxymethylene carbon (-OCH-), and methyl carbon (-CH_3_) signals typically occur in the region of δ 170, δ 69, and δ 17, respectively, thus representing the configurational structure of PLA. As shown in Fig. 4C, the molecular weight of PLA was determined by GPC and MALDI-TOF mass spectroscopy (Fig. S3) ^12^. From the GPC spectrum analysis, the average molecular weight (M_w_) was found to be 62.5 kDa, and the molecular weight distribution ranged from 2 kDa to 100 kDa (Fig. S2). The result is approximately consistent with the MALDI-TOF mass spectrum (Fig. S3).

**Fig. 3.**
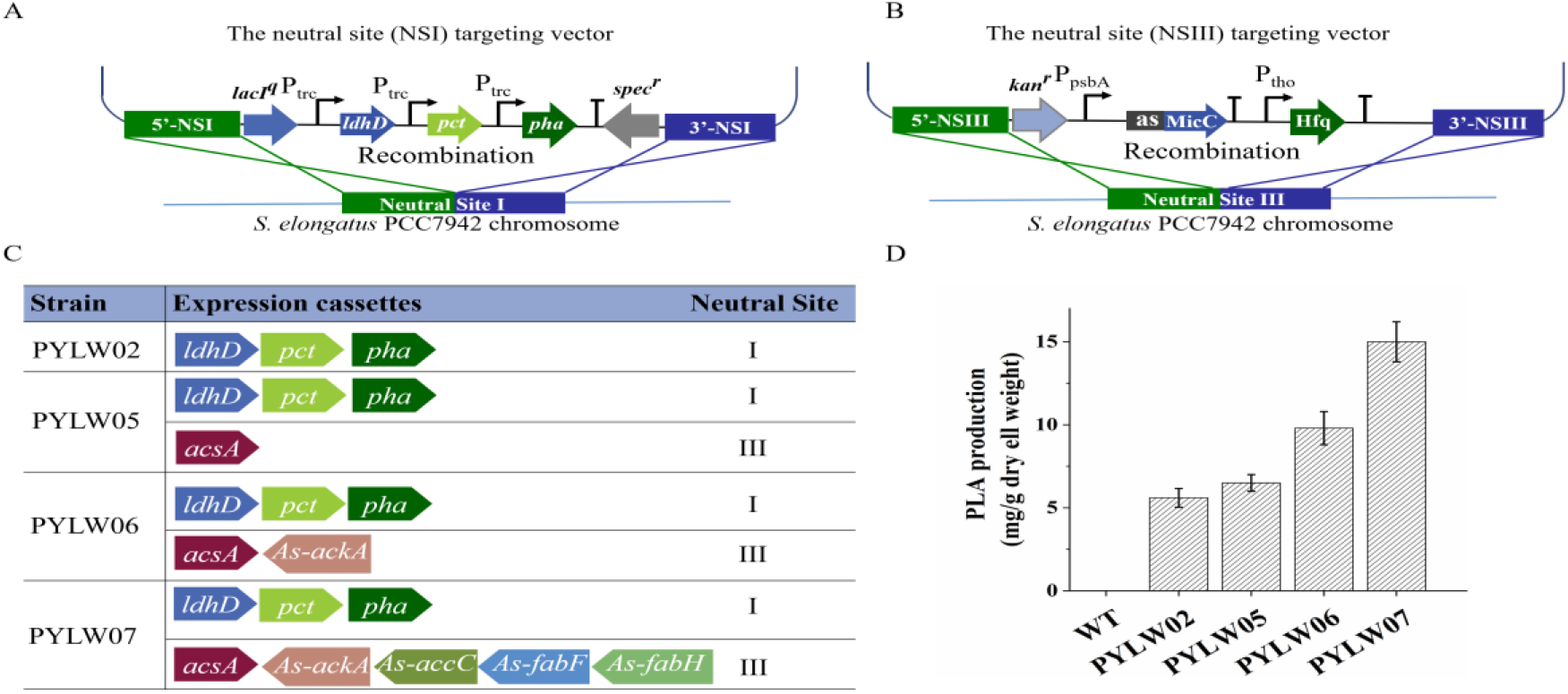
The production of PLA by engineered *S. elongatus* PCC7942 using different expression cassettes. (A) Schematic representation of inserting the PLA expression cassettes into the *S. elongatus* PCC7942 chromosome at the NSI. (B) The schematic representation of inserting the sRNA expression cassettes into *S. elongatus* PCC7942 chromosome at the NSIII. (C) List of engineered strains with different recombination. The positive arrows represented the overexpressed genes and the arrows in reverse indicate the genes were knocked down by the sRNA tool. (D) The PLA production of the different recombination strains. Error bars indicate SD (n = 3)

**Fig. 4.**
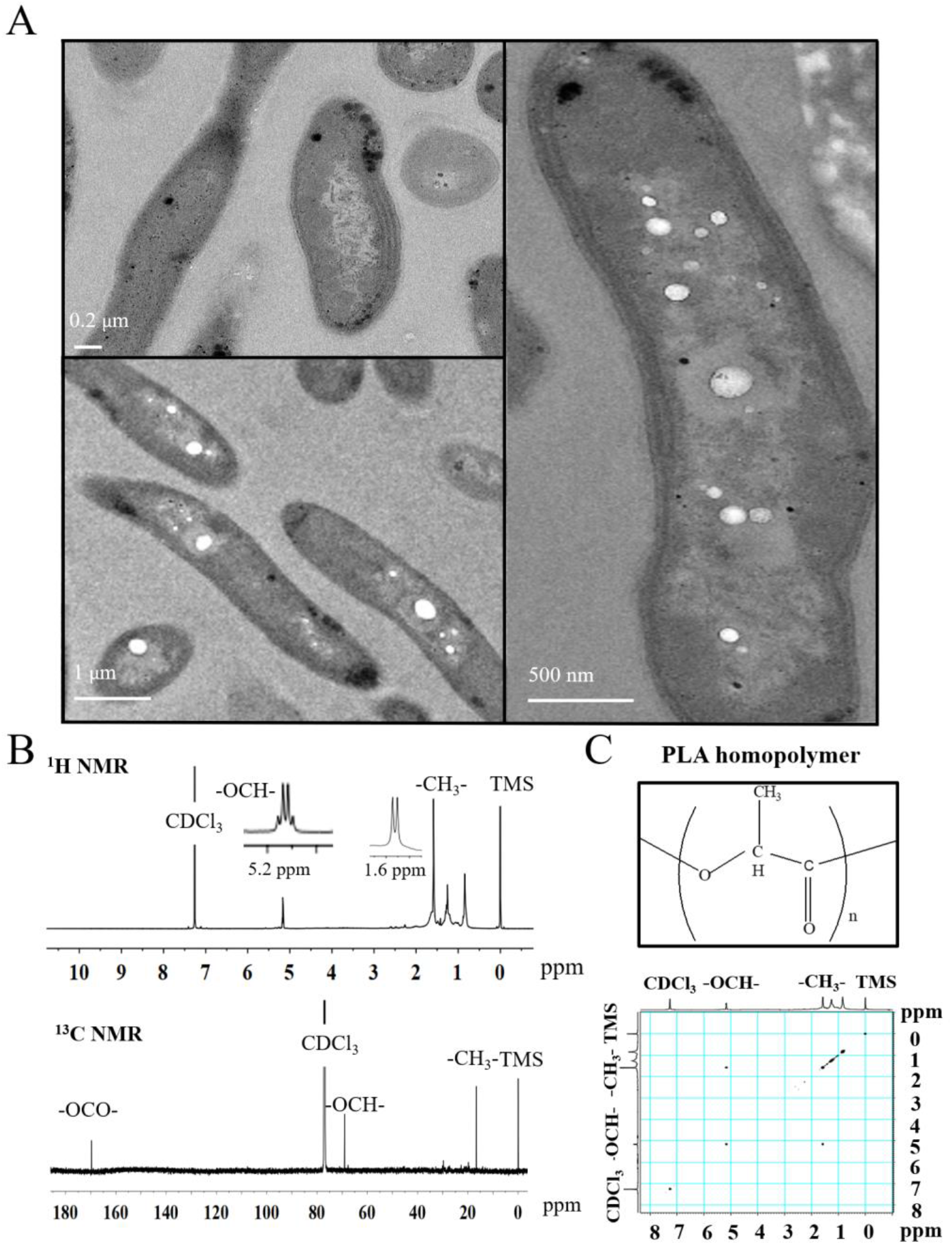
The characteristics of PLA homopolymer accumulated inside of recombinant *S. elongatus* PCC7942 cells. (A) The transmission electron micrographs of intracellular PLA homopolymer: Top left panel, the wild type of *S. elongatus* PCC7942 cells. The right and bottom left panel, the PLA homopolymer granules accumulated in recombinant *S. elongatus* PCC7942 cells. (B) The ^1^H, ^13^C and COSY NMR spectra of PLA homopolymer produced by the recombinant *S. elongatus* PCC7942 cells. The right box showed the structure of the PLA homopolymer.

### Promoter optimization for the PLA biosynthesis pathway

The recombinant *S. elongatus* harboring the metabolic engineering pathway for PLA biosynthesis could produce PLA, but at very low efficiency, yielding 0.8 mg/g DCW PLA (Fig. 2B). Based on the previous study, together with our study ^7,13^, the rate-limitation steps of PLA biosynthesis are believed to be the generation of lactyl-CoA and the polymerization step. Thus, the expression of PCT and PHA synthase was optimized under the control of P_trc_, P_psbA_ and P_cpc560_, compared with the three genes under the control of one promoter. The quantitative reverse transcription PCR (RT-qPCR) was then performed to test the transcriptional levels of the two heterologous genes of strain PYLW01. As shown in Fig. 2D, the genes *pct* and *pha* were all successfully transcribed under the different promoters. It was found that the transcriptional level of the two rate-limiting genes was the highest under control of promoter P_trc_, out of the four other constructs. Furthermore, the four different recombinants were cultivated under P_trc_ in the BG11 medium, to induce the heterologous enzymes, with constant light exposure to produce PLA. Subsequently, the highest level of production was achieved under the control of three independent promoters of P_trc_. The yield of the resulting strain PYLW02 increased approximately 7-fold to 5.6 mg/g DCW PLA (Fig. 2B). It is in accordance with RT-qPCR data. Thus, it appears that the P_trc_ promoter is the most effective for producing PLA, judging from both the expression levels and the yield data. Therefore, the strain PYLW02 was used for the subsequent experiments.

### Designing of an acetyl-CoA self-sufficient system to increase CoA pool

Based on the *in silico* genome-scale metabolic flux analysis done in the previous study ^26^, the surplus acetyl-CoA pool was expected to increase PLA production, since it might be one of the bottlenecks for PLA synthesis. To further enlarge the acetyl-CoA pool and enhance PLA production, we constructed an acetyl-CoA self-sufficient system in the strain PYLW02. Firstly, the native promoter of *acsA* encoding the acetyl-CoA synthetase was replaced with the stronger P_trc_ promoter. Then, the P_trc_-*acsA* expression cassettes were integrated into the neutral site III, resulting in the strain PYLW05 (Fig. 3C). The sRNA regulatory tool has been successfully applied to the cyanobacteria to knock down four genes (with up to 96% blockage), while hardly imposing any metabolic burden on host cells ^14,15^.

Moreover, considering that the gene *ackA* encoding acetate kinase can convert acetyl-CoA to acetate, we knocked down *ackA* using sRNA, resulting in the strain PYLW06 (Fig. 3C). The PLA production slightly increased after overexpression of acetyl-CoA synthetase, and was significantly improved upon knocking down the gene *ackA*; the yield of the strain PYLW06 increased approximately 12-fold to 9.8 mg/g DCW, compared with the strain PYLW01(Fig. 3D). These results indicate that the biosynthesis efficiency of PLA was enhanced by increasing the amount of acetyl-CoA pool.

### Redirection of carbon flux to the PLA biosynthetic pathway

As the fatty acid biosynthesis could compete for the carbon flux with the biosynthesis of bioproducts ^16^, it is necessary to regulate the expression of crucial fatty acid pathway enzymes, so that more carbon flux is directed to PLA biosynthesis. In this study, sRNA was also used to limit the amount of acetyl-CoA channeling to fatty acid synthesis ^17^ to increase PLA production further (Fig. 3). In cyanobacteria, FabH converting acetyl-CoA to acetyl-ACP, FabF converting acetyl-ACP to acetoacetyl-ACP, and ACC (acetyl-CoA carboxylase, encoding by *accABCD* gene) converting acetyl-CoA to malonyl-CoA, have been shown to be essential for fatty acid biosynthesis (Fig. 1) ^18,19^. Previously, the downregulation of the critical genes *fabD, fabB, fabF*, and *fabH*, successfully decreased fatty acid production, thus increasing malonyl-CoA levels up to 5-fold using PTRNAs ^16^. In this study, *accC, fabF*, and *fabH*, were downregulated to redirect the carbon flux to the PLA biosynthesis pathway using sRNA, as shown in Fig. 1 and Fig. 3. The resulting strain PYLW07 was cultivated in BG11 medium for PLA production, after being induced by IPTG and theophylline, resulting in approximately 19-fold increase to 15 mg/g DCW, compared with the strain PYLW01(Fig. 3D).

## Discussion

Disposable plastic items are ubiquitous worldwide today. Unfortunately, most used plastics produced from fossil resources are non-degradable, causing serious environmental issues. Therefore, degradable plastics will inevitably take the place of regular plastics with time ^20^. Meanwhile, atmospheric CO_2_ levels have increased sharply in the past 50 years to approximately 414 ppm, after staying stable around 200∼280 ppm for around 40,000 years ^9,21^. It may significantly contribute to global warming, rise in sea levels, and even the extinction of ∼24% of plant and animal species ^2^. It is a worldwide consensus that effective strategies for carbon capture need to be developed ^22,23^ to meet the climate-change severe mitigation challenge and adhere to the Paris Agreement of 2015.

Furthermore, PDLA is a vital feedstock to improve the thermal resistance, mechanical performance, and hydrolysis resistance of PLA based materials ^24^. Poly (L-lactic acid) (PLLA) and PDLA can form stereocomplex (SC) crystals with a high melting point, that can be used to synthesize the high-performance of PLA ^25^. In addition, no metal catalyst is involved in the biosynthesis of PDLA, which makes it suitable for use in pharmaceutical and medical industries ^26,27^. Surprisingly, the highest reported average molecular weight for *in vivo* PDLA production (M_w_, 62.5 kDa, M_n_, 32.8 kDa, and dispersity of 1.9) was obtained in this work (Fig. S2, Fig. S3). In accordance with previous studies, the highest melting peak was obtained when the number-average molecular weight (M_n_) of PDLA ranged from 23 to 50 kDa ^28^. The M_n_ of the PDLA produced by cyanobacteria was in the optimal range, which will facilitate its use for the synthesis of the high-performance of PLA and endow extra commercial value on the products.

Currently, the production of PLA is mainly carried out by the chemical polymerization of lactic acid, which is primarily produced by the microbial fermentation of sugars. Unfortunately, the production of lactic acid from the sugar-based feedstocks unavoidably competes with global food supplies ^29^. Compared with the sugar-based feedstock (called first-generation feedstock) and food-related biomass (called second generation feedstock), CO_2_ (called the third generation feedstock) is increasingly desirable for bioproduction processes ^9^. Therefore, the use of CO_2_ for the production of PLA appears to be more attractive.

## Methods

### Chemicals and reagents

Unless otherwise specified, all chemicals were purchased from Sigma-Aldrich (St. Louis, MO) or Macklin (GmbH, Germany). The restriction enzymes, T4 DNA ligase (New England Biolabs Inc, USA), 2 × Phanta Max Master Mix (Vazyme, Nanjing, China), were used for cloning and plasmid construction. The polyacrylamide flocculant and polyelectrolyte flocculant were purchased from the Jieyuan Chemical Co., LTD (Cangzhou, China). Genomic DNA was extracted using the Wizard Genomic DNA Purification kit (Promega, Madison, WI, USA) and other reagents of high quality were obtained from general suppliers in Shanghai.

### DNA manipulation

All DNA manipulation and general molecular biology techniques were performed according to standard protocols ^30^. The plasmids and primers used in this study were listed in Table S2, Table S3 and briefly described. The pEASY-Blunt cloning vector (Transgen, China) was used for subcloning genes. The shuttle vector pAM-MCS12 was used for inserting the gene cassettes into the neutral site I (NS I) of the *S. elongatus* PCC7942 chromosome ^17^. The Hfq-MicC expressing plasmid, inserting the small RNA tools into the neutral site III (NS III) of the *S. elongatus* PCC7942 genome, was a kind gift from Dr. Tao Sun ^31^. The *ldhD* gene encoding D-lactate dehydrogenase was engineered to switch the coenzyme specificity from NADH to NADPH, which has been done before ^7^, thus promoting NADPH utilization in cyanobacteria for D-lactate production. The genes encoding propionyl-CoA transferase (PCT) from *Clostridium propionicum* and the PHA synthase from *Pseudomonas* sp. were synthesized by GenScript (Nanjing, China) with codon optimization based on the codon bias of *Synechococcus elongates* ^10^. The gene *acsA* encoding acetyl-CoA synthetase was from the native genome of the *S. elongatus* PCC7942. Plasmid pAM-PYLW01 was constructed by inserting *ldhD, pct* and *pha* into pAM-MCS12 plasmid ^32^ between the *Eco*RI and *Xho*I sites under the control of the IPTG-inducible promoter P_trc_. Thereafter, the plasmid pAM-PYLW02, pAM-PYLW03, pAM-PYLW04 were generated by adding the independent P_trc_ promoter, P_psbA_ promote and P_cpc560_ promoter to targeting *pct* and *pha* gene integrated at NSI, respectively. In addition, the strain of PYLW05 was generated by insertion of the *acsA* into the NSIII site using the pBA3031M plasmid. Moreover, knockdown of the *ackA* (acetate kinase), *accC* (acetyl-CoA carboxylase), *fabF* (ketosynthase), *fabH* (β-Ketoacyl-ACP Synthase III) genes was performed using the inducible Hfq-MicC system integrated at NSIII ^31^, generating the strain of PYLW06 and PYLW07.

### Strain construction and transformation

The strains used in this study are listed in Table S1. All the operations of chromosomes were carried out by homologous recombination of the plasmid DNA into the *S. elongatus* PCC7942 chromosomes at the neutral site I or neutral site III ^33^. The strain of PYLW01, PYLW02, PYLW03, PYLW04 were constructed by recombination of plasmids pAM-PYLW01, pAM-PYLW02, pAM-PYLW03 and pAM-PYLW04 into the chromosomes of *S. elongatus* PCC7942, respectively. In addition, the strain PYLW05 was then constructed by recombination of plasmid DNA of pBA-*acsA* into the strain PYLW02 chromosomes at neutral site III. Moreover, the expression cassettes of sRNA were introduced into the chromosomes at neutral site III using the Hfq-MicC tool. The P_psbA_ promoter was used for expressing all sRNAs and the chaperone *hfq* was under the control of a theophylline-responsive riboswitch. The gene of *ackA, accC, fabF, fabH* were downregulated by sRNA tools. Schematic of the PYLW06 and PYLW07 strain were listed in Fig. 3C. *S. elongatus* PCC7942 was transformed using previously described protocols ^34^. All the transformants were cultivated in the BG11 medium with the 20 mg/L spectinomycin for successful selection recombination. The right candidate transformants were further analyzed by PCR using the primers listed in Table S3 to verify that the targeted genes were correctly integrated into the NSI or NSIII of the *S. elongatus* PCC7942.

### PLA extraction and analysis

The PLA-produced culture was centrifuged at 8,000 rpm for 15 min, and then the cells were washed twice with distilled water and freeze-dried overnight at -80 °C. The PLA was extracted using the soxhlet apparatus with chloroform ^35^. About 2.0 g lyophilized cells were filled with 200 volumes of the chloroform refluxed in the soxhlet apparatus for 24 h, and the extracted materials were collected. The chloroform-polymer mixed solution was filtered. The polymer was washed twice in the ten volumes of hexane and collected by centrifugation ^36^.

GC-MS determined the concentration of the polymers accumulated in the cells. The cultures were centrifuged at 8,000 rpm for 15 min, and then the cells were washed twice with distilled water and freeze-dried overnight at -80 °C. About 60 mg of samples were dissolved in acidified methanol, i.e., containing 15% (v/v) H_2_SO_4_ and chloroform in a hydrothermal synthesis reactor at 120 °C for 4 h. The resulting methyl esters of constituent lactate were subsequently used for the GC-QQQMS analysis ^11,37^. The GC-MS analysis was conducted by Thermo & TSQ8000 system equipped with TraceGOLD TG-WAXMS (30 m × 0.25 mm×0.25 μm). The GC oven temperature was initially at 50 °C for 5 min and ramped to 230 °C at 7.5 °C/min. Then the oven temperature was increased to 260 °C with a gradient of 10 °C/min and held for 5 min.

The composition of the polymer was determined by NMR on a Bruker AVANCE III HD 600 MHz (Bruker, Rheinstetten, Germany) using the tetramethylsilane (TMS) as an internal standard. The polymer was diluted in CDCl_3_ and the NMR spectra were recorded at 298 k. The ^1^H NMR spectrum was acquired using an excitation flip angle of 30°at the delay time of 10 s. In the ^13^C NMR experiment, the 30°pulse width was 3.5 μs at the delay time of 2.5 s. The 2D COSY experiment was performed with 1.3 s delay time, 16 scans and processing data size of 2 k × 1 k.

The molecular weights of the polymer were determined by gel permeation chromatography (GPC) at 35 °C using the waters1525 system equipped with waters 2414 detectors and Agilent PLgel Sum MIXED-C column. The elution solvent used was chloroform at a flow rate of 1 mL/min. Samples were resuspended in chloroform and filtered through a 0.4 μm filter. The MALDI-TOF mass spectrum was carried out using the DCTB as a matrix, CF_3_CO_2_Na as a cationization agent and chloroform as a solvent in a MALDI-7090 (SHIMADZU, Japan) mass spectrometer ^12^.

### Transmission electron microscopy analysis

The wild type and engineered *S. elongatus* PCC7942 cells were fixed with glutaraldehyde and then post-fixed with potassium permanganate for 1 h. Then, the cells were dehydrated in graded ethanol and propylene oxide, respectively. The Leica EM UC7 system conducted the ultrathin sections after being stained with lead citrate and uranyl acetate ^38^. The sample was then examined using the biological transmission electron microscopy (Tecnai G2 spirit Biotwin system) at 80 kHz.

### RT-qPCR analysis of synthetic pathway genes

The engineered *S. elongatus* PCC7942 strains were grown in BG11 medium at 30 °C with an illumination intensity of 100 mol photons m^−2^ s^−1^ when the OD_730_ reached 0.4-0.6, 1 mM IPTG was added to the culture to induce the targeted genes. The total RNA was then extracted using an RNAprep Pure Cell/Bacteria Kit (Tiangen Biotech, China). The extracted total RNA was then treated with DNase I to remove the genomic DNA and used as a template for cDNA synthesis with random primers and SuperScriptTMIII Reverse Transcriptase (Invitrogen, China). The quantitative PCR was conducted using the SuperReal PreMix SYBR Green kit (Tiangen Biotech, China) and the CFX96 Real-Time system (Bio-Rad, Hercules, CA, USA). Beacon Designer was used to designing the primers for qPCR and the *rnpB* gene was used as a control.

## Acknowledgements

This work was supported by the grants of National Key R&D Program of China (2018YFA0903600) and NSFC (31870088 and 32170105). We thank Dr. Tao Sun for his kind gift of sRNA plasmid.

## Author contributions

Fei Tao and Ping Xu conceived the study. Chunlin Tan performed the experiments and analyzed the data. Chunlin Tan wrote the manuscript. Fei Tao and Ping Xu critically reviewed the manuscript and revised it. All authors read and approved the submitted version.

## References

1 de Albuquerque, T. L., Marques Junior, J. E., de Queiroz, L. P., Ricardo, A. D. S. & Rocha, M. V. P. Polylactic acid production from biotechnological routes: A review. 186, 933–951, International journal of biological macromolecules, https://10.1016/j.ijbiomac.2021.07.074 (2021).

2 Hughes, T. P. et al. Climate change, human impacts, and the resilience of coral reefs. Science 301, 929–933, https://10.1126/science.1085046 (2003).

3 Yang, T. H. et al. Tailor-made type II Pseudomonas PHA synthases and their use for the biosynthesis of polylactic acid and its copolymer in recombinant Escherichia coli. Applied microbiology and biotechnology 90, 603–614, https://10.1007/s00253-010-3077-2(2011).

4 Singhvi, M. S., Zinjarde, S. S. & Gokhale, D. V. Polylactic acid: synthesis and biomedical applications. Journal of applied microbiology 127, 1612–1626, https://10.1111/jam.14290(2019).

5 Li, G. et al. Synthesis and biological application of polylactic acid. Molecules 25, 5023–5041, https://ARTN502310.3390/molecules25215023 (2020).

6 Lajus, S. et al. Engineering the yeast yarrowia lipolytica for production of polylactic acid homopolymer. Frontiers in bioengineering and biotechnology 8, 954–969, https://ARTN95410.3389/fbioe.2020.00954 (2020).

7 Li, C. et al. Enhancing the light-driven production of D-lactate by engineering cyanobacterium using a combinational strategy. Scientific Reports 5, 9777–9788, https://ARTN0977710.1038/srep09777 (2015).

8 Okano, K. et al. Efficient production of optically pure D-lactic acid from raw corn starch by using a genetically modified L-lactate dehydrogenase gene– deficient and alpha–amylase–secreting lactobacillus plantarum strain. Applied and environmental microbiology75, 462–467, https://10.1128/Aem.01514-08(2009).

9 Liu, Z. H., Wang, K., Chen, Y., Tan, T. W. & Nielsen, J. Third–generation biorefineries as the means to produce fuels and chemicals from CO_2_. Nature catalysis 3, 274–288, https://10.1038/s41929-019-0421-5(2020).

10 Yang, T. H. et al. Biosynthesis of polylactic acid and its copolymers using evolved propionate CoA transferase and PHA synthase. Biotechnology and bioengineering 105, 150–160, https://10.1002/bit.22547(2010).

11 Jung, Y. K., Kim, T. Y., Park, S. J. & Lee, S. Y. Metabolic engineering of Escherichia coli for the production of polylactic acid and its copolymers. Biotechnology and bioengineering 105, 161–171, https://10.1002/bit.22548(2010).

12 Chen, C. J. et al. ppm–Level Thermally switchable yttrium phenoxide catalysts for moisture-insensitive and controllably immortal polymerization of rac-lactide. Macromolecules 51, 6800–6809, https://10.1021/acs.macromol.8b01229(2018).

13 Matsumoto, K. et al. Improved production of poly(lactic acid)-like polyester based on metabolite analysis to address the rate-limiting step. Amb express 4, 83– 88, https://10.1186/s13568-014-0083-2(2014).

14 Gaida, S. M., Al-Hinai, M. A., Indurthi, D. C., Nicolaou, S. A. & Papoutsakis, E. T. Synthetic tolerance: three noncoding small RNAs, DsrA, ArcZ and RprA, acting supra–additively against acid stress. Nucleic acids research 41, 8726– 8737, https://10.1093/nar/gkt651(2013).

15 Sun, T. et al. Toolboxes for cyanobacteria: Recent advances and future direction. Biotechnology advances 36, 1293–1307, https://10.1016/j.biotechadv.2018.04.007(2018).

16 Yang, Y., Lin, Y., Li, L., Linhardt, R. J. & Yan, Y. Regulating malonyl-CoA metabolism via synthetic antisense RNAs for enhanced biosynthesis of natural products. Metabolic engineering 29, 217–226, https://10.1016/j.ymben.2015.03.018(2015).

17 Ni, J., Tao, F., Wang, Y., Yao, F. & Xu, P. A photoautotrophic platform for the sustainable production of valuable plant natural products from CO_2_. Green chemistry 18, 3537–3548, https://10.1039/c6gc00317f(2016).

18 Parsons, J. B. & Rock, C. Bacterial lipids: Metabolism and membrane homeostasis. Progress in lipid research 52, 249–276, https://10.1016/j.plipres.2013.02.002(2013).

19 Kuo, J. & Khosla, C. The initiation ketosynthase (FabH) is the sole rate-limiting enzyme of the fatty acid synthase of Synechococcus sp PCC 7002. Metabolic engineering 22, 53–59, https://10.1016/j.ymben.2013.12.008(2014).

20 Choi, S. Y. et al. Metabolic engineering for the synthesis of polyesters: A 100-year journey from polyhydroxyalkanoates to non-natural microbial polyesters. Metabolic engineering 58, 47–81, https://10.1016/j.ymben.2019.05.009(2020).

21 Petit, J. R. et al. Climate and atmospheric history of the past 420,000 years from the vostok ice core, Antarctica. Nature 399, 429–436, https://Doi10.1038/20859 (1999).

22 Pikaar, I. et al. Carbon emission avoidance and capture by producing in–reactor microbial biomass based food, feed and slow release fertilizer: Potentials and limitations. Science of the total environment 644, 1525–1530, https://10.1016/j.scitotenv.2018.07.089(2018).

23 Mac Dowell, N., Fennell, P. S., Shah, N. & Maitland, G. C. The role of CO_2_capture and utilization in mitigating climate change. Nature climate change 7, 243–249, https://10.1038/Nclimate3231(2017).

24 Han, X. et al. Steps Toward high-performance PLA: economical production of D-lactate enabled by a newly isolated Sporolactobacillus terrae strain. Biotechnology journal 14, 1088656–1088667, https://ARTN180065610.1002/biot.201800656 (2019).

25 Lv, T. X. et al. New insight into the mechanism of enhanced crystallization of PLA in PLLA/PDLA mixture. Journal of applied polymer science 135, 45663– 45670, https://ARTN4566310.1002/app.45663 (2018).

26 Jozwicka, J., Gzyra–Jagiela, K., Gutowska, A., Twarowska–Schmidt, K. & Cieplinski, M. Chemical purity of PLA fibres for medical devices. Fibres & textiles in eastern europe 20, 135–141 (2012).

27 Ata, R. et al. Recent applications of polylactic acid in pharmaceutical and medical industries. Journal of chemical and pharmaceutical research 2015, 51–63 (2015).

28 Shao, J. et al. Remarkable melting behavior of PLA stereocomplex in linear PLLA/PDLA blends. Industrial & engineering chemistry research 54, 2246– 2253, https://10.1021/ie504484b(2015).

29 Varman, A. M., Yu, Y., You, L. & Tang, Y. J. J. Photoautotrophic production of D–lactic acid in an engineered cyanobacterium. Microbial cell factories 12, 117– 125, https://Artn11710.1186/1475-2859-12-117 (2013).

## References

30 Chong, L. Molecular cloning -A laboratory manual, 3rd edition. Science 292, 446–446, https:/10.1126/science.1060677(2001).

31 Sun, T. et al. Re-direction of carbon flux to key precursor malonyl-CoA via artificial small RNAs in photosynthetic Synechocystis sp. PCC 6803. Biotechnology biofuels 11, 26–42, https://10.1186/s13068–018–1032–0(2018).

32 Wang, Y., Tao, F., Ni, J., Li, C. & Xu, P. Production of C3 platform chemicals from CO_2_by genetically engineered cyanobacteria. Green chemistry 17, 3100– 3110, https://10.1039/c5gc00129c(2015).

33 Bustos, S. A. & Golden, S. S. Expression of the psbDII gene in Synechococcus sp. strain PCC 7942 requires sequences downstream of the transcription start site. Journal of bacteriology 173, 7525–7533, https://10.1128/jb.173.23.7525–7533.1991 (1991).

34 Golden, S. S., Brusslan, J. & Haselkorn, R. Genetic engineering of the cyanobacterial chromosome. Methods enzymology 153, 215–231, https://10.1016/0076-6879(87)53055-5(1987).

35 Jung, Y. K. & Lee, S. Y. Efficient production of polylactic acid and its copolymers by metabolically engineered Escherichia coli. Journal of biotechnology 151, 94–101, https://10.1016/j.jbiotec.2010.11.009(2011).

36 Nomura, C. T., Taguchi, K., Taguchi, S. & Doi, Y. Coexpression of genetically engineered 3–ketoacyl–ACP synthase III (fabH) and polyhydroxyalkanoate synthase (phaC) genes leads to shor-chain-length-medium-chain-length polyhydroxyalkanoate copolymer production from glucose in Escherichia coli JM109. Applied and environmental microbiology 70, 999–1007, https://10.1128/AEM.70.2.999-1007.2004(2004).

37 Braunegg, G., Sonnleitner, B. & Lafferty, R. M. A rapid gas chromatographic method for the determination of poly-β-hydroxybutyric acid in microbial biomass. European journal of applied microbiology and biotechnology 6, 29–37, https://10.1007/BF00500854(1978).

38 Koch, M. et al. Maximizing PHB content in Synechocystis sp. PCC 6803: a new metabolic engineering strategy based on the regulator PirC. Microbial cell factories 19, 231–243, https://ARTN23110.1186/s12934-020-01491-1 (2020).

